# Heme minimizes Parkinson’s disease-associated toxicity by inducing a conformational distortion in the oligomers of alpha-Synuclein

**DOI:** 10.1101/629238

**Authors:** Ritobrita Chakraborty, Sandip Dey, Simanta Sarani Paul, Pallabi Sil, Jayati Sengupta, Krishnananda Chattopadhyay

## Abstract

Aggregation of the intrinsically disordered protein alpha-Synuclein (α-Syn) into insoluble fibrils with a cross-β sheet amyloid structure plays a key role in the neuronal pathology of Parkinson’s disease (PD). The fibrillation pathway of α-Syn encompasses a multitude of transient oligomeric forms differing in size, secondary structure, hydrophobic exposure and toxicity. According to a recent solid state NMR study, the fibrillating unit of α-Syn contains the core residues of the protein arranged into in-register parallel β sheets with a unique Greek key topology. Here, we have shown that the physiologically available small molecule heme (hemin chloride) when added at sub-stoichiometric ratios to either monomeric or aggregated α-Syn, arrests its aggregation in an oligomeric state, which is minimally toxic. Using cryo-EM, we observed that these heme-induced oligomers are ‘mace’-shaped and consist of approximately four monomers. However, the presence of a crucial twist or contortion in their Greek key structural architecture prevents further hierarchical appending into annular oligomers and protofilament formation. We confirm using a His50Gln mutant that the binding of heme onto His50 is crucial in inflicting the structural distortion and is responsible for the stabilization of the non-toxic and off-pathway α-Syn oligomers. We believe that this study provides a novel strategy of developing a therapeutic solution of PD, which has been elusive so far.

## Introduction

A plethora of evidences associate the fibrillation of the intrinsically unfolded protein alpha synuclein (α-Syn) with the neurodegenerative movement disorder Parkinson’s disease (PD) (1, 2). PD is characterised by the presence of fibrillar intracellular Lewy body plaques within the dopaminergic neurons of the substantia nigra of the mid-brain. *In vitro*, α-Syn has been shown to follow the nucleation-conversion-polymerization model of aggregation (3, 4). The kinetics of this pathway initiates with the primary nucleation phase, in which the native monomeric protein misfolds to form nuclei which combine and subsequently elongate to give rise to different classes of oligomers, protofilaments and fibrils. Current hypotheses predict that the initial oligomeric intermediates of fibrillar structures, common to many neurodegenerative disease-related proteins including α-Syn, are the primary toxic species. Upon maturation, the fibrils which are otherwise minimally toxic (5), fragment into oligomers which act as secondary nuclei or ‘seeds’ for *de novo* fibrillation giving rise to a feedback loop of fibril amplification (6).

Several structural descriptions of mature fibrillar assemblies manifest the universality of the amyloid cross-β architecture (7). A solid state NMR study has revealed a ‘Greek-key’ β-sheet topology of a fibrillar assembly of α-Syn monomers (8), which also coincides with the conserved ‘bent β arch kernel’ architecture of both the rod and twister fibril polymorphs observed recently (9). Although 3D architectures of α-Syn fibril assembly determined by high-resolution cryo-EM, have been published very recently (9–11), structural understanding of the intermediate species which precedes fibrillation is limited. Some structural evidences of oligomeric structures describe annular intermediates with a central pore, which might explain the characteristic pore-forming activity (12) of α-Syn protofibrils during membrane permeabilization (6, 13). A more recent study has identified lipid binding structural elements at the N-terminal of a specific oligomeric species of α-syn that are responsible for cell disruption and consequently give rise to cellular toxicity (14). As a result, structural insights into the heterogeneous classes of oligomers formed in the early stages of the aggregation pathway of α-syn have been deemed as necessary in order to design potential therapeutic strategies against PD.

Heme has been reported to interact with α-Syn leading to the formation of soluble oligomers (15), suggesting that this may be the mechanism by which filament formation is inhibited. Additionally, ferric dehydroporphyrin IX and related macrocyclic compounds can inhibit amyloid fibril formation with IC_50_ values in the low micromolar range (16–18). Under physiological conditions within the human brain and peripheral RBCs, the heme-containing neuronal hemoglobin scavenges α-Syn, leading to a reduction of PD-induced mitochondrial damage and apoptosis (19). Additionally, over-expression of heme-containing neuroglobin inside neuronal cells, which also express α-Syn, reduces cytoplasmic α-Syn inclusions and associated mitochondrial damage (20). These results encouraged us to obtain a deeper insight into the mechanism underlying the prevention of fibrillation of α-Syn upon the addition of heme.

The addition of heme (in the form of hemin chloride) to either monomeric α-Syn (at 0 h, preincubation) as well as to a heterogeneous population of prefibrillar and fibrillar aggregates (after 48 h of aggregation, late incubation) at a sub-stoichiometric molar ratio yielded small morphologically similar oligomeric structures, suggesting that these structures are the stable endpoints of the heme-induced inhibition of aggregation. We employed cryo-electron microscopy (cryo-EM) for the structural characterization of these similar-sized heme-treated oligomeric forms which revealed that heme primarily stabilizes a ‘mace’-shaped oligomer comprising approximately four monomers. This structure is presumed to be the fundamental unit of the fibrillar aggregates of α-Syn as it closely resembles the recently proposed solid state NMR-based ‘Greek key’ model of the elementary unit of a protofilament of α-Syn (8, 9). Nevertheless, closer analysis of the molecular architecture of the heme-stabilized mace oligomer indicated that the structure represents a distorted version of the Greek key. We therefore termed the heme-treated oligomer with the distorted Greek key motif as the ‘twisted Greek key oligomer’ (8, 9).

Recent advances in the 3D structure elucidation of the fibrils of α-Syn using cryo-EM have demonstrated that the monomeric units folded into the Greek key motif stack upon each other with their longitudinal axes parallel to each other, to form a protofilament. Two such protofilaments intertwine down the length of their longitudinal axes along a tightly packed steric zipper interface to form either the mature rod-like or twister-like fibril polymorphs (9). The conformational distortion in the twisted Greek key oligomers occurs upon heme binding to the His50 residue of α-Syn located at (rod polymorph) or near (twister polymorph) the inter-protofilament hydrophobic steric zipper interface, which weakens/ prevents the binding between the two protofilaments, thereby disintegrating the fibril (9). The role of His50 was confirmed by employing a His50Gln (H50Q) mutant, which did not offer heme-induced inhibition of aggregation. Additionally, using FT-IR, we observed a reduction in the antiparallel β sheet component of the oligomers that were treated with heme. In accordance to previous reports (18, 21, 22), this finding can be correlated with the toxicity of the heme-treated twisted Greek key oligomeric population that showed reduced transcellular seeding propensity as well as low pathogenicity to both biomembrane-mimicking liposomes and neuroblastoma cells, as compared to the on-pathway fibril-forming oligomers. Our results have allowed us to propose the molecular mechanism of heme-mediated inhibition of α-Syn fibrillation. The identification and characterization of the twisted Greek key oligomer as the fundamental unit of α-Syn amyloid assembly provides insights into the mechanism of fibril formation and sheds light on opportunities for therapeutic intervention at various stages of aberrant protein self-assembly.

## Results and Discussion

### Heme inhibits the primary and secondary nucleation micro-events within the fibrillation pathway of α-Syn

We used Thioflavin T (ThT) fluorescence to investigate the effect of heme on the different microscopic events that occur within the amyloid assembly process of α-Syn. In the absence of heme, we observed the distinctive sigmoidal aggregation kinetics (23, 24) of α-Syn (Figure 1A). The ThT data was complimented by direct imaging by negative stain TEM and AFM, which clearly showed the formation of a fibrillar network of α-Syn (Figure 1B). The diameter of the mature fibrils after 96 hours of incubation in the absence of heme typically varied between 8.5-10 nm (TEM data, Figure 1B), while the mean height measured 9.7 nm (AFM data, Figure 1B). The length of the fibrils measured between 0.5-2 μm. Upon addition of heme at the beginning of the aggregation reaction (pre-incubation; α-Syn/heme 25:1), the protein showed insignificant ThT fluorescence, indicating the heme-induced inhibition of the primary nucleation-elongation micro-events in a dose-dependent manner (Figure 1A). This was supported by the TEM and AFM results which showed that upon pre-incubation with heme, fibril formation did not occur even after 96 hours of incubation under aggregation-inducing conditions (Figure 1C). Instead, the aggregation was stopped at the oligomer stage, the diameters of which varied between 1 and 3.5 nm, while the mean height according to AFM measurements was 2.3 nm (Figure 1C). These oligomers formed when α-Syn is treated with heme from the beginning of the aggregation period have been named oligomers_1_ in this study.

**Figure 1:**
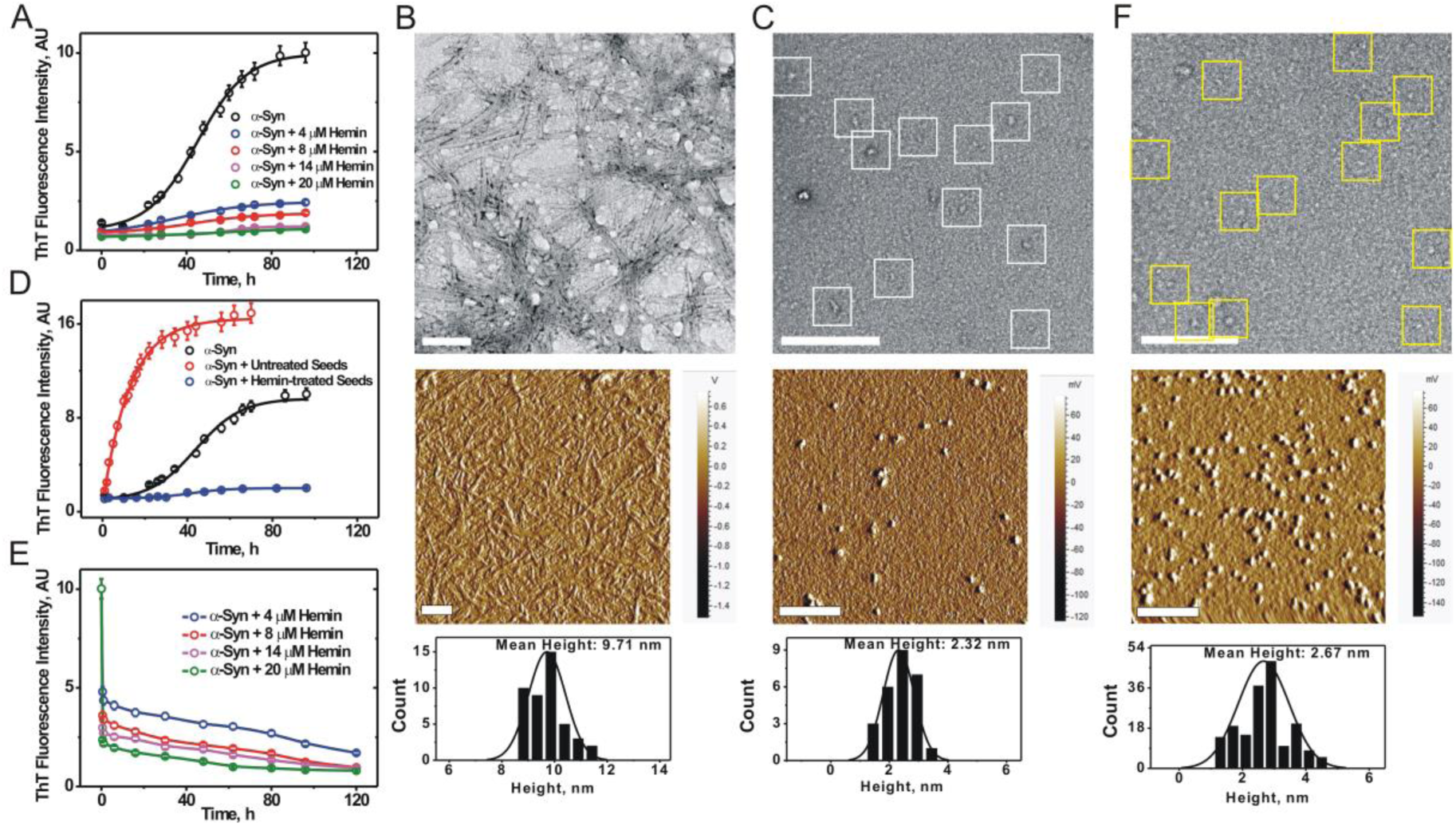
Heme arrests the aggregation of α-Syn by converting the aggresome into oligomers. (A) Dose-dependent study (using ThT fluorescence) of 200 μM monomeric α-Syn pre-incubated with heme. (B-C) TEM and AFM micrographs of (B) α-Syn (96 h); α-Syn oligomers_1_ (white squares) formed in presence of heme (pre-incubation; α-Syn/heme 25:1). (D) Comparison of the seeding effect of heme-treated (α-Syn/heme 25:1, in blue) and untreated seeds (in red) on monomeric α-Syn; the aggregation kinetics profile of 200 μM α-Syn (in black) has been added to this figure for comparison. (E) Dose-dependent study (using ThT fluorescence) of the disaggregation of 96 h fibrils, upon addition of heme (α-Syn/heme 25:1). (F) TEM and AFM micrographs depicting the formation of oligomers_2_ (yellow squares) upon disaggregation by heme of 96 h fibrils (α-Syn/heme 25:1). The scale bars in the TEM and AFM micrographs are 100 nm and 200 nm, respectively. The corresponding height distribution histograms are presented beneath each AFM image.

In contrast to the usual sigmoidal behaviour, the presence of pre-existing fibril seeds resulted in hyperbolic aggregation kinetics (Figure 1D). However, seeds prepared from fibrils that had been incubated in presence of heme appeared to be inert and completely inhibited seed-induced aggregation (Figure 1D). We subsequently investigated the effect of heme on the processes of fibril fragmentation and secondary nucleation within the amyloid amplification cycle. When heme was added to the fibrillar aggregates that are formed after 96 h of incubation under aggregation-inducing conditions, (α-Syn/heme 25:1, late incubation) it caused a dose-dependent and stable reduction in ThT fluorescence (Figure 1E), suggesting the inhibition of secondary nucleation on preformed fibril surfaces. The TEM image (Figure 1F) depicting the breakdown of the 96 h aggregates showed an aggresome that predominantly comprised a homogeneous population of spherical oligomers upon incubation with heme (α-Syn/heme 25:1). The mean height of 2.7 nm of these oligomers was calculated from the corresponding AFM image (Figure 1F). Collectively, these results demonstrate that heme can arrest the multi-pathway fibrillation processes and convert the heterogeneous aggresome into a population of predominant oligomeric species, which shall be henceforth termed as oligomers_2_.

### Heme reduces seeding capacity and toxicity of α-Syn oligomers_1_ and oligomers_2_

Within the brain of patients suffering from PD, the on-pathway α-Syn aggregates spread transcellularly along neural networks and seed *de novo* aggregation via prion-like mechanisms (25, 26). To demonstrate the off-pathway (i.e. non-fibril forming) non-seeding nature of the heme-stabilised oligomers, *in vitro* seeding of exogenously-added heme-treated as well as untreated fibril seeds was studied in SH-SY5Y cells that were transiently transfected with an EGFP-α-Syn construct. Figure 2A shows the confocal image of cells transfected with the GFP construct that were subsequently transduced with 1 μM sonicated fibril seeds for 24 h. At this concentration, the majority of cells displayed punctate intracellular inclusions. Importantly, cells treated with fibrils prepared in presence of heme (Figure 2B) or those transfected with α-Syn-GFP but not transduced with any seeds (vehicle, Figure 2C) did not convert to an inclusion-positive state.

**Figure 2:**
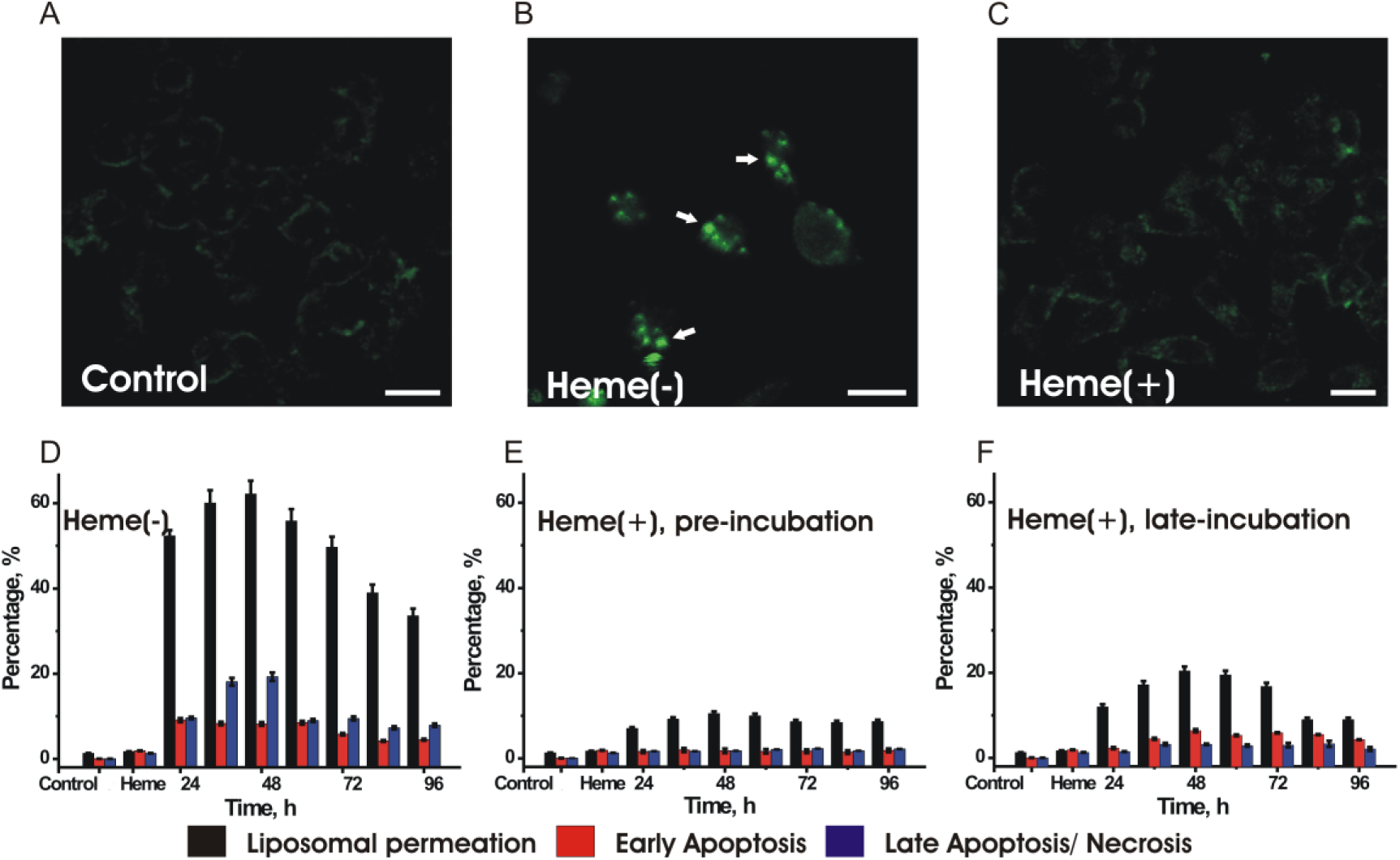
Heme minimizes *in cell* seeding, synthetic lipid membrane permeabilization and cytotoxicity of various aggregated species of α-Syn. (A-C) Exogenously added fibril seeds prepared from recombinant protein, induce endogenous α-Syn aggregation in α-Syn-GFP-transfected SH-SY5Y cells. Transfected cells were transduced with (A) 0 μM (control) or 1 μM seeds and incubated for 24 h. Seeds were prepared by sonicating pre-formed α-Syn fibrils that had been incubated under aggregation-inducing conditions for 96 h either in the (B) absence or (C) presence of heme. Punctate green structures within the cytoplasm in (B) denote seeded α-Syn aggregates. The scale bars denote 20 μm. Effect of α-Syn species formed at various time-points (D) incubated without heme; (E) pre-incubated with heme from the beginning of aggregation period; (F) incubated with heme at various stages of aggregation, on calcein-containing liposomal SUV permeation; early apoptosis and late apoptosis/ necrosis as observed in SH-SY5Y neuroblastoma cells.

Subsequently, we compared the toxicity of the various structural forms of the protein that are formed during the aggregation pathway in the absence and presence of heme. For this purpose, we collected aliquots from the aggregation reaction of the protein after every 12 hours. In the first assay, the species formed in the absence (on-pathway oligomers, Figure 2D) and presence of heme (Figure 2E: oligomers_1_, pre-incubation; 2F: oligomers_2_, late incubation) were treated with calcein-loaded 3: 7 POPC: DOPS small unilamellar vesicles (SUVs) at a protein: lipid ratio of 1:10. Pore formation induced by the different structural forms of α-Syn lead to permeable vesicles releasing calcein, the extent of which indicated the toxicity of that particular protein form. Additionally, we quantified the percentage of SH-SY5Y neurobastoma cells undergoing early and late apoptosis/ necrosis in presence of the heme-treated or untreated structural species of α-Syn. In accordance to previous reports (13), we observed that the heterogeneous population of oligomers, protofilaments and protofibrils, (these structures were observed from AFM measurements, Figure S1A, SI) formed between 24 h and 48 h of incubation induce maximum toxicity in both SUV membranes as well as neuroblastoma cells. The heme-stabilised structures (pre-incubation with heme: oligomers_1_ or late incubation with heme: oligomers_2_) at the equivalent time-points showed significantly reduced toxicity (Figure 2E and 2F, respectively).

### Structural characterization of the heme-stabilized oligomers

A previous study (13) has described using cryo-EM that the fibril-forming on-pathway oligomers formed after 20-24 h of aggregation are annular. These oligomers possess a high content of anti-parallel β sheets and show liposomal permeability as well as cytoxicity. In order to study the mechanism of heme-induced prevention of fibrillation of α-Syn, we prepared these on-pathway oligomers and compared their structure with that of the heme-pre-incubated oligomers_1_ that were formed by incubating the monomeric protein with heme (α-Syn/Heme 25:1) under aggregation-inducing conditions for 24 h. Additionally, we also studied the structure of the oligomers_2_ that were formed when the heterogeneous aggresome formed after 48 h of aggregation was treated with heme (late incubation).

Using FPLC-size exclusion chromatography, we isolated the oligomers_1_ and oligomers_2_, both of which eluted at an identical retention time of 17.1 minutes. This proves that the addition of heme for both pre-incubation or late incubation conditions populates oligomers of identical molecular weight/ size (Figure S1B, SI). FT-IR measurements were then used to compare the secondary structures (27) of the untreated on-pathway oligomers with the purified heme treated oligomers_1_ and oligomers_2_. The on-pathway oligomers formed after 24 h of incubation in the absence of heme that showed augmented toxicity, contained a large extent (12%) of antiparallel β sheet structure (Figure 3A) evidenced by component bands at ∼1685-95 cm^-1^ along with four times prominent bands at 1620-38 cm^-1^, in accordance with previous studies (13). In contrast, the oligomers_1_ (Figure 3B) and oligomers_2_ (Figure 3C) contained a much reduced antiparallel β sheet component (∼2%). Similar β sheet orientation in the oligomeric species of α-Syn have been reported previously (28), and oligomer populations with differing characteristics have been shown to cause difference in the membrane perturbation and cell death via apoptosis (18, 21, 22). The low frequency bands at ∼1617 cm^-1^ are associated with β-strands in aggregated structures indicating inter-molecular β sheet structure with strong hydrogen bonds.

**Figure 3:**
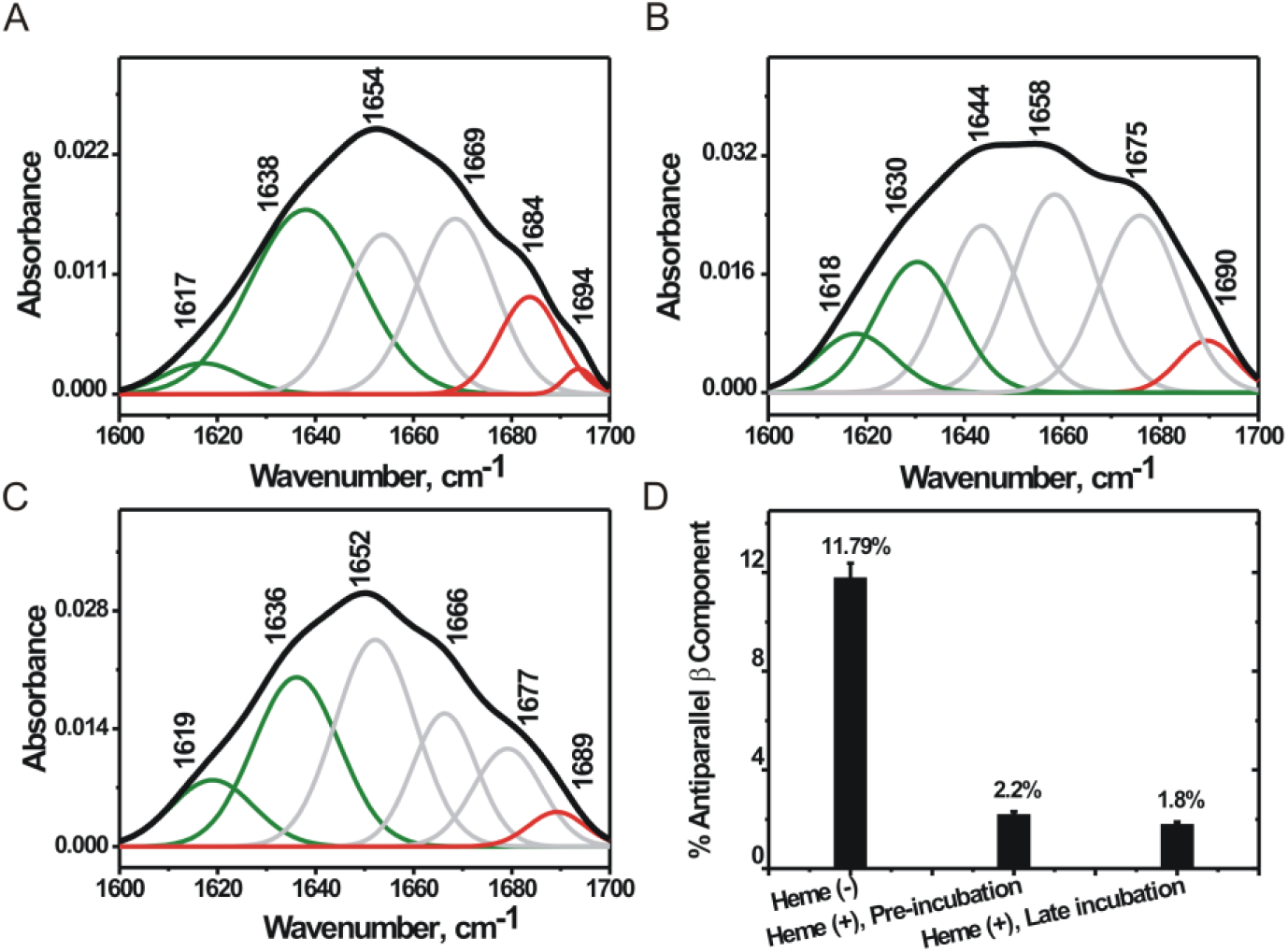
Heme causes a major reduction in the antiparallel β sheet component of fibril-forming aggregates. Deconvoluted FT-IR spectra of 24 h oligomers incubated (A) in the absence of heme; or (B) in presence of heme from the beginning of incubation at 0 h (oligomers_1_, pre-incubation); (C) depicts the IR spectrum of oligomers_2_ formed when 48 h prefibrillar and fibrillar aggregates were treated with heme. (D) Addition of heme causes a drastic reduction in the antiparallel component (bands at ∼1680-90 cm^-1^, Figure A-C, illustrated in red) of the resulting oligomeric population which may be correlated with the decrease in cytotoxicity. The bands depicted in green denote the parallel β sheet component (1622-1638 cm^-1^).

Subsequently, we used cryo-EM in conjunction with single particle reconstruction technique (29) to characterize the structures of heme-treated α-Syn oligomers_1_ (dataset 1; early treatment) and oligomers_2_ (dataset 2, late incubation) formed in the presence of heme. Negative stain TEM imaging clearly showed (Figure 1C & F) the formation of small oligomers following heme treatment at either pre-incubation or late incubation stage. The cryo-EM micrographs of dataset 1 (Figure 4A) revealed predominant distribution of small particles of uniform shape and size (∼ 6-8 nm). Particle picking, and initial model building were performed using EMAN2 (30). Reference-free two-dimensional (2D) classifications of the particles were performed in different image processing programs (Figure 4B: I, II & III). Final reference-based (using the EMAN2 initial model) 3D reconstruction and refinement were done in SPIDER (31) (Figure 4C). To ensure the reliability of the map, we used several validation strategies (see Methods and Supplementary Figure S2B, S2D and S2F).

**Figure 4:**
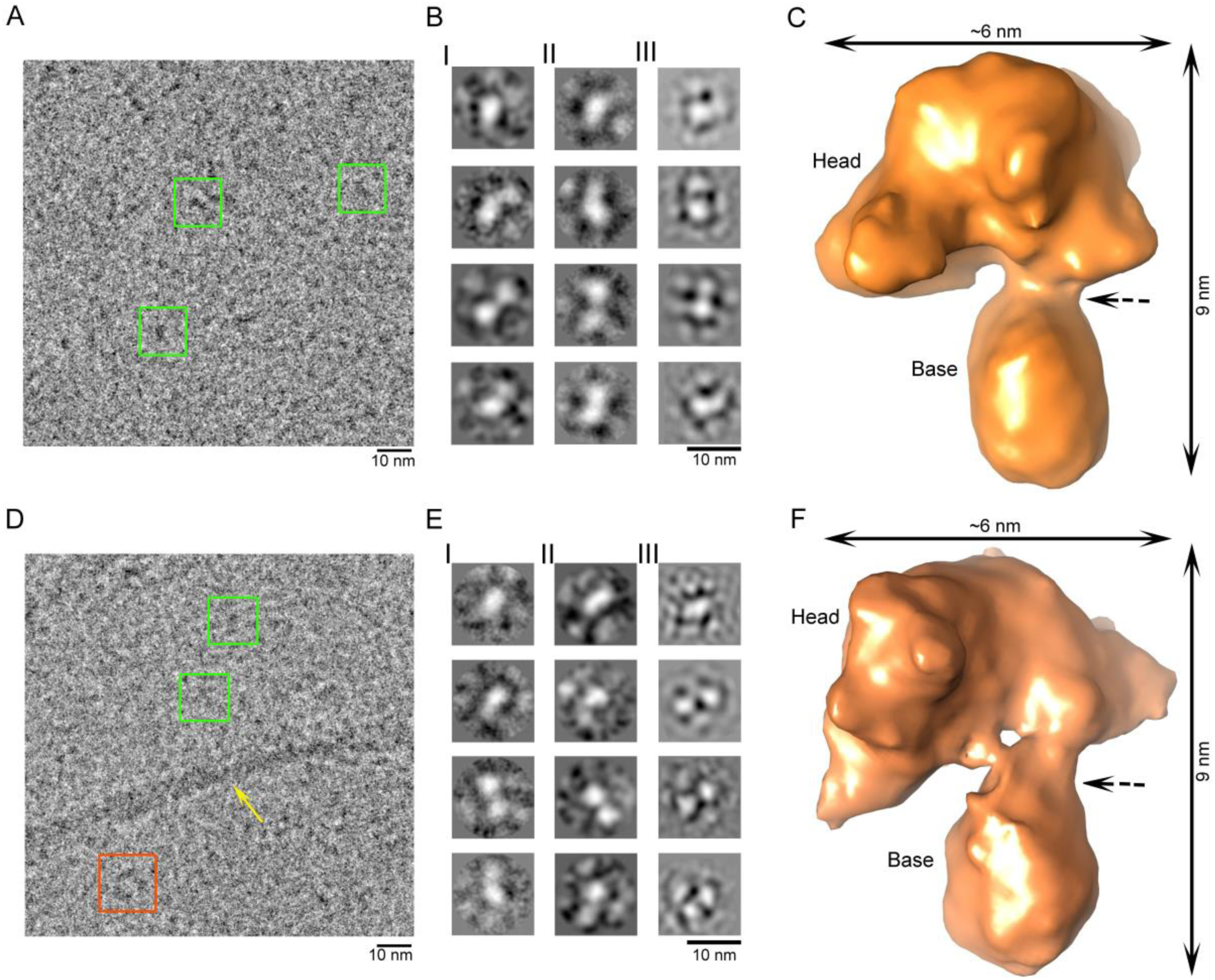
Cryo-EM study of heme-stabilized α-Syn oligomers. (A) Raw micrograph showing uniformly distributed small oligomers (green box, oligomers_1_). (B I-III) Reference free 2D class averaging analysis were done by Xmipp, RELION and EMAN2. (C) Cryo-EM density map of α-Syn oligomers_1_. (D-E) Heme treatment was done after fibril formation, i.e., after 48 h of aggregation. (D) Raw micrograph showing distribution of small oligomers (oligomers_2_) similar to oligomers_1_, annular oligomers and fibrils were marked by green box, orange box and yellow arrow, respectively. (C I-III) Reference free 2D class averaging analysis were done by Xmipp, RELION and EMAN2. (F) Cryo-EM density map of α-Syn oligomers_2_. (C-F) Dotted black arrows indicate the kinks between head and base of both oligomers.

The cryo-EM micrographs of dataset 2 in contrast, showed an ensemble of particles of different sizes along with few short length fibrillar structures (Figure 4D). We selected the population of oligomers (oligomers_2_) that appeared qualitatively similar in size to the particles seen for oligomers_1_ (∼6-8 nm; Figure 4D). These small size particles were selectively picked, following which three reference-free 2D analyses were done (Figure 4E), and a 3D cryo-EM map was generated following the same procedure used for oligomers_1_ (Figure 4F). Both the 3D cryo-EM maps (Figure 4C and 4F) appeared as a club with a heavy head (‘mace oligomer’). The resolutions of the maps generated from dataset 1 and dataset 2 were estimated to be ∼12.6 Å and 12 Å, respectively, as determined by the 0.5 FSC criterion (Figure S2A and S2C).

Thus, it is evident that the mace oligomer is the fundamental structural unit formed whenever heme is added to α-Syn at any stage (early or late) in its aggregation pathway. Earlier literatures have reported α-Syn oligomers to be annular in shape of different dimensions (13, 32). We have also identified the formation of a horseshoe-shaped oligomer when α-Syn was allowed to aggregate for 20-24 h in the absence of heme (Figure S2, S3) which appeared to be a precursor of the annular forms (Figure S3C). Interestingly, the presence of mace-like units could be detected within the structure of the on-pathway horseshoe oligomer formed in the absence of heme (Figure S3D, E), indicating that the mace-like units probably assemble to form the annular oligomers. We propose that the individual mace oligomers are unstable and hence immediately assemble to form higher order annular structures. However, heme stabilizes the mace oligomers and also disintegrates the annular forms (or protofibrils/ mature fibrils) into their constituent mace oligomers.

### Heme stabilizes the ‘molecular mace’ architecture by interacting with His50

Our results suggest that the assembly of more than one mace oligomers of α-Syn leads to the formation of an annular oligomer. However, heme stabilizes the mace oligomer in such a manner that its propagation into higher order oligomers and fibrillar structures is terminated. In order to comprehend the molecular mechanism underlying the stabilization of the mace oligomers by heme, we measured the binding affinity of heme to the tetramethylrhodamine-5-maleimide (TMR)-tagged monomeric α-Syn Gly132Cys (G132C) mutant, by studying the heme-induced quenching of the dye. Since wild type α-Syn does not have a cysteine residue, a G132C mutation was introduced for the maleimide labelling. This single cysteine mutant G132C is considered similar to the wild type protein for biophysical studies (23, 33). While the dissociation constant (K_d_) and number of binding sites on the α-Syn G132C monomer for heme was 590 nM and 1 respectively (Figure 5A), heme binding to α-Syn was not observed when excess (10 mM) imidazole was present (Figure S4A, SI), suggesting a probable binding of the heme to a histidine residue in the protein.

**Figure 5:**
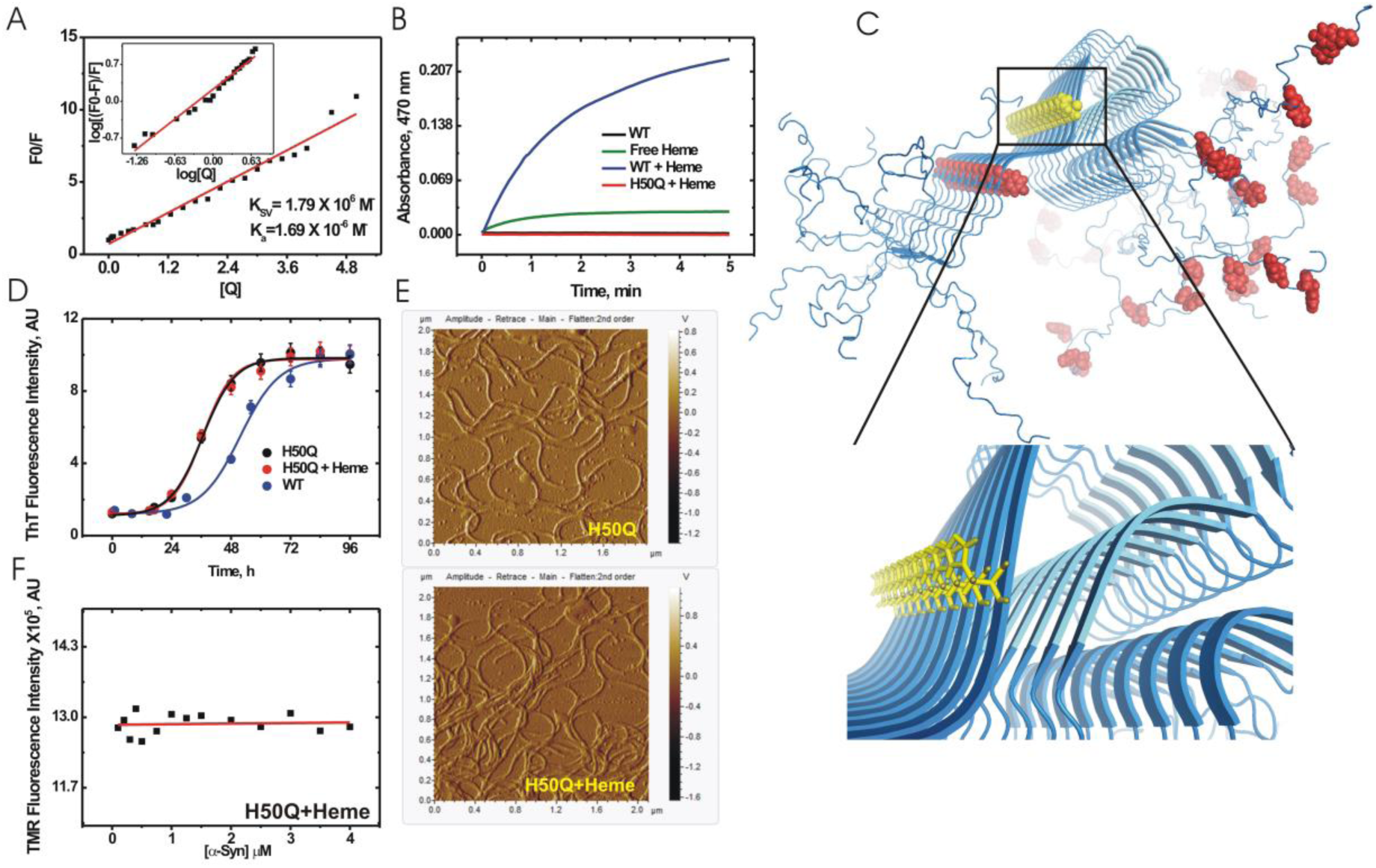
Heme binds to the His50 residue within the NAC region of the Greek key oligomer of α-Syn forming a peroxidase-positive complex. (a) Stern-Volmer plot and double log plot (inset) obtained from the steady-state quenching and binding of TMR-tagged α-SynWT with heme. (b) Kinetic traces for peroxidase activity monitored for the α-SynWT-heme complex. (c) Greek key β-sheet alignment of the fibril core with the potential heme-binding residues (Tyr: in blue; His: in red). Inset shows magnified view. (d) The aggregation behaviour of the histidine mutant H50Q is unaltered in presence of heme as observed using ThT fluorescence. (e) AFM micrographs depict that H50Q aggregates into fibrils even in the presence of heme. (f) The H50Q mutant shows no binding with heme, as observed from the absence of any quenching of TMR-tagged H50Q by heme.

The heme-α-Syn WT complex was also found to have a considerably high peroxidase (His-coordinated heme enzyme) activity (34) compared to hemin chloride (Figure 5B). These experiments further indicated the involvement of His residues in the heme-α-Syn binding. A molecular modelling exercise of the heme-induced mace oligomer structure suggested His50, Tyr39, Tyr125, Tyr133 and Tyr136 as five potential residues, which are present on the solvent-exposed surface of the mace structure and could be involved in the oligomer-oligomer or oligomer-heme stabilization (Figure 5C). Since His50 is the only histidine positioned at this region, we prepared a His50Gln (H50Q) mutant. Although H50Q aggregated to an extent similar to the WT protein, its aggregation was not inhibited by the addition of heme (Figure 5D: ThT; 5E: AFM). Moreover, the H50Q mutant did not show any binding to heme as observed from the fluorescence binding measurement (Figure 5F) and had no peroxidase activity in presence of heme either (Figure 5B). These results confirmed the crucial role of His50 in the heme-mediated stabilization of the mace structure. Incidentally, H50Q is a missense mutation that causes a late-onset familial form of PD and dementia (35). We propose that the absence in this mutant, of the only His residue that can bind to the physiologically-prevalent heme and thus salvage fibrillation is responsible for its susceptibility to aggregation.

As mentioned previously, a recent NMR study identified a unique conformation of an oligomeric unit (9) (comprising ten monomers) of the α-Syn fibril with a Greek-key motif (8). More recently, cryo-EM studies revealed similar Greek-key folds / ‘kernels’ in α-Syn fibrils (9, 10). Our oligomers_1_ and oligomers_2_ had a SEC retention time of 17.1 minutes that correspond to a MW of ∼65 kDa (Figure S1C), providing preliminary evidence that these oligomers contain approximately 4 monomeric units (MW of monomeric α-Syn 14.5 kDa). Both oligomers_1_ oligomers_2_ show striking resemblances with the recently described Greek-key structural motif consisting of ∼ four monomers (Figure 6A) (36). The Greek-key architecture fits well in the C-terminal or head region of the molecular mace structure. However, a distortion was observed at the junction of the head and the base of the heme-stabilized mace oligomer giving rise to the ‘twisted Greek key oligomer’ (Figure 6B-C). Segmentation (in Chimera) of the horseshoe-shaped oligomeric intermediate formed in the absence of heme resulted in three segments of similar shape and size. Interestingly, a tetrameric unit of α-Syn with the Greek key motif could be accommodated well into each of the segments in Chimera (Figure S3D, E). Thus, apparently, tetrameric units containing the Greek key motif are initially formed which subsequently self-assemble to form annular structures of different dimensions. We hypothesize that heme binding to the His50 residues on the exposed surface of the tetrameric Greek key structure either at (rod fibril polymorph) or near (twister polymorph) the head-base junction (Figure 7) induces a torsion (Figure 4C and 4F) and thereby dislodges further amyloid assembly.

**Figure 6:**
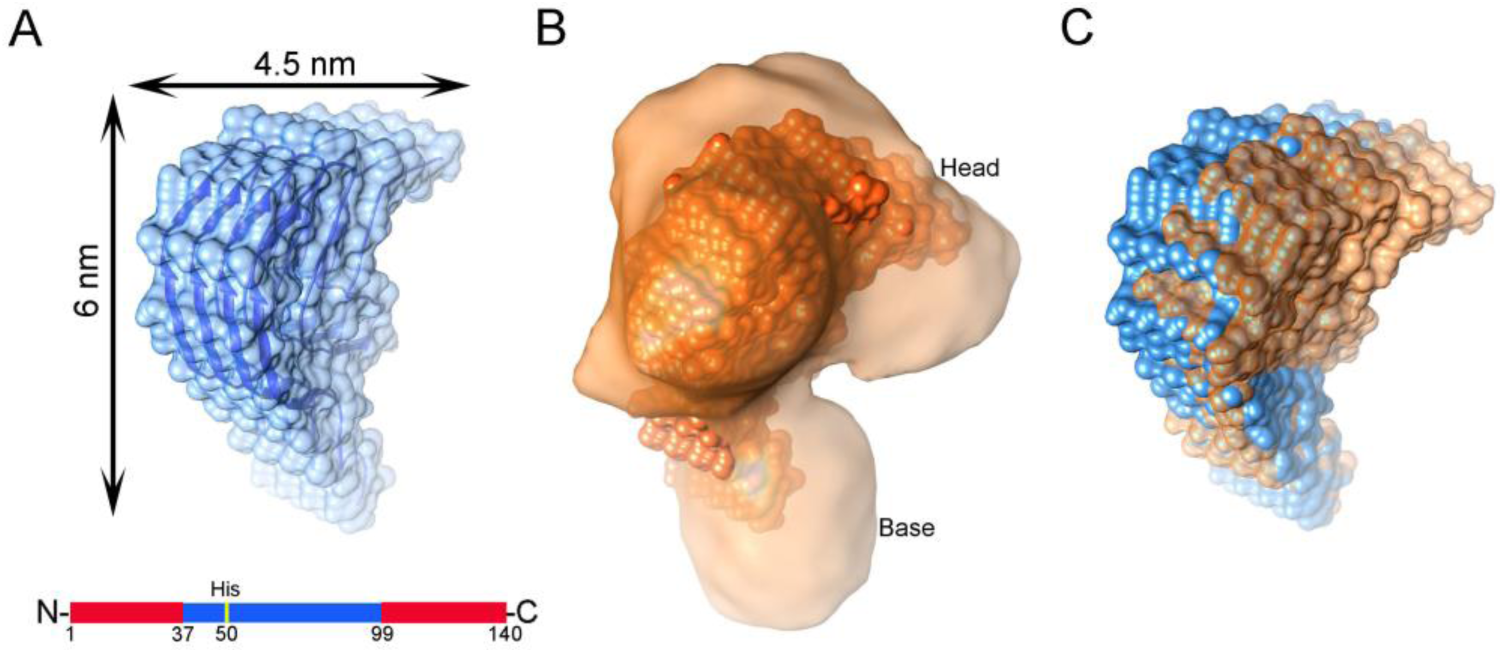
Structural characterization of the molecular mace density map. (A) View of the greek-key structure (2N0A) consisting of 4 monomeric chains in cartoon representation (blue) inside the semitransparent surface. Only the structured part (residue 37-99) is shown and the unstructured regions (residues 1-36 and 100-140, shown below) are not included in the structure. (B) Full structure shown in (A) could not be fitted into the mace density as a rigid piece. The ‘head’ and ‘base’ parts of the Greek key motif (containing 4 monomers) was fitted separately as a rigid body (sphere representations (CPK) in orange) into the molecular mace density map generated from dataset 1 (semi-transparent orange). (C) Superimposition of the greek key structure (blue) and the fitted model (orange), represented in spheres, showed distortion in the fitted model.

**Figure 7:**
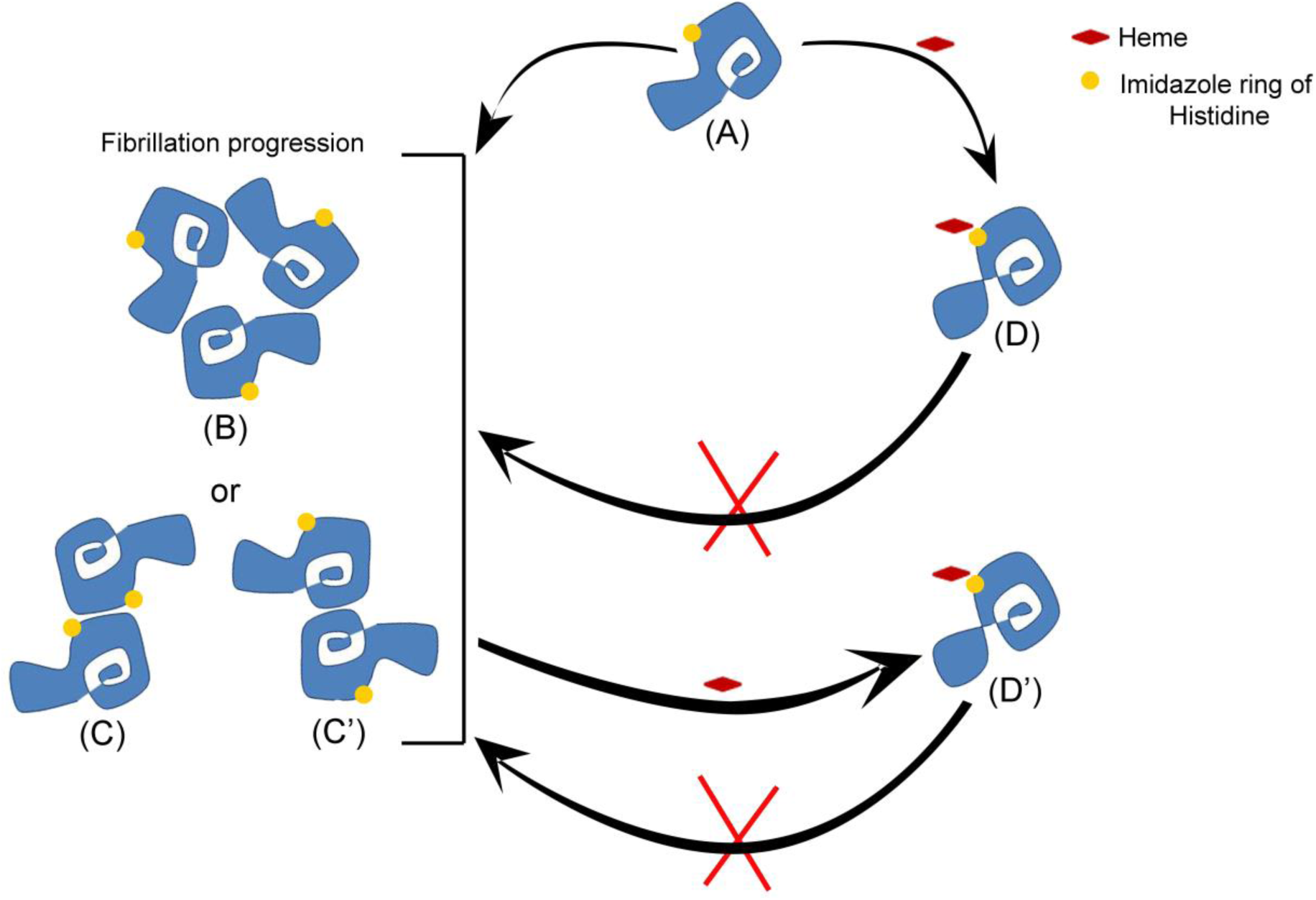
Schematic representation of the proposed mechanism of heme-mediated inhibition of α-Syn fibrillation. Our hypothesis entails that the perturbation of the native structure of α-Syn (38, 39) leads to the formation of an on-pathway oligomer having the β sheet Greek key motif (8), which is presumably the fundamental unit of fibrillation (A). Fibril elongation may occure either via formation of intermediate oligomers which eventually assemble to form fibrils (B), or via initial stacking of Greek key oligomers into protofibrils which subsequently entwine to form fibrils (C or C’). having either rod or twister polymorphs (30). Addition of heme at an initial stage arrests the fundamental oligomeric unit by stabilizing it into a twisted form (D) which is unable to assemble and form fibrillar structures. Addition of heme at a later stage (when fibrillation has already initiated) causes disassembly of α-Syn oligomers/fibrils by inducing a twist/ distortion in the fundamental oligomeric units resulting in a oligomeric form with a twisted Greek key motif (D’) that appears very similar to (D).

Thus, our structural analyses along with biochemical results suggested that α-Syn oligomerization proceeds via beta-stacked structure containing the Greek key motif as proposed by Tuttle et al (8). Upon heme treatment, either during progression of fibrillation or degradation of mature fibrillar structures, heme prefers to bind to an oligomeric form comprising ∼4 associated monomers. Heme binding distorts this structure into a ‘twisted Greek key’ resulting in an inert ‘off pathway’ oligomer that shows reduced cytotoxicity. As a consequence of this distortion, further association of monomers to form mature amyloid fibrils is inhibited (Figure 7).

## Conclusion

By studying the mechanism of heme-induced fibril breakdown, we propose the probable hierarchical fibril construction machinery of α-Syn. Fibril assembly may occur by the initial formation of annular oligomers, each of which are composed of Greek key oligomers joined end-to-end (Figure 7A-B and S3D). These annular oligomers then stack on each other to form mature fibrils as has been reported earlier (13). Additionally, fibril assembly can also transpire via the initial formation of protofibrils (Figure 7C & C′), each composed of Greek key oligomers stacked with their longitudinal axes parallel to each other (37). These protofibrils (diameter ∼5 nm) then intertwine along their longitudinal axes along a steric zipper interface to form mature fibrils having a diameter of ∼10 nm. Figure 7C & C′ depict the cross-section of the rod (Figure 7C) and the twister fibril polymorphs (Figure 7C′) showing the location of the heme-binding His50 residue. However, the amalgamation of the Greek key oligomers into either annular oligomers or elongated protofibrils is halted in the presence of heme through one of the following two pathways. In the first pathway, heme binding to the His50 residues of the Greek key oligomer causes a torsion at its head-neck interface (Figure 7D) leading to the formation of the twisted Greek key oligomer. This arrests further association into either annular oligomers or protofibrils. In the second mechanism, preformed annular oligomers and protofilaments break down into the twisted version of the Greek key oligomer upon heme-binding, thereby compromising the binding interfaces and preventing further fibrillation (Figure 7D′).

The second mode of heme-mediated inhibition of fibrillation involves the hydrophobic preNAC steric zipper geometry (i.e., residues Gly51-Ala56) of the fibril rod polymorph. The protofilament interface is stabilized by a salt bridge between His50 and Glu57′ (Glu57′ belongs to the opposite protofilament). We propose that the heme competes with Glu57′ in binding to the His50 residue, thus destabilizing the zipper interface and disintegrating the fibril into its two constituent protofilaments. Stabilization of such a non-toxic off-pathway oligomeric form of α-Syn would be a potential pharmacological approach for PD treatment.

## Materials and Methods

For the purification of α-Syn, Tris salt, isopropyl β-D-galactopyranoside (IPTG), phenylmethanesulfonylfluoride (PMSF), ammonium sulfate, urea, sodium dihydrogen sulfate, disodium hydrogen sulfate and sodium chloride from Sigma-Aldrich (MO, USA) were used. Thioflavin T, calcein and hemin chloride were purchased from Sigma-Aldrich. Glycerol and sodium dodecyl sulfate (SDS) were purchased from Merck (NJ, USA) and USB (OH, USA), respectively. For high performance liquid chromatography (HPLC), two columns from Waters Corporation (MA, USA) were used. For the liposome permeation study, the lipids were obtained from Avanti Polar Lipids (AL, USA). FBS, DMEM, Opti-MEM were obtained from Life Technologies (Carlsbad, CA). Cell culture media, supplements, kits and reagents were purchased from Invitrogen, CA, USA, unless specified otherwise. The dye Tetramethylrhodamine-5-maleimide was purchased from Invitrogen (CA, USA). All other chemicals used for this study were obtained in the highest grade available.

### Expression and purification of α-Syn protein

Recombinant human α-Syn wild type (WT), G132C and H50Q/G132C mutants were expressed in *Escherichia coli* BL21 (DE3) strain transformed with the pRK172 α-Syn wild type or mutant plasmid. All site-directed mutagenesis were performed using the Agilent Quikchange Lightning site-directed mutagenesis kit (Agilent Technologies, CA, USA). The expression was induced by adding 1 mM IPTG to a culture that had reached an OD of 0.5-0.6. The cultures were then incubated at 37 °C with shaking at 180 rpm for 4 hours. Cells were harvested by centrifugation. The cell pellets were then resuspended in sonication buffer (10 mM Tris, pH 7.4 and 1 mM PMSF) and lysed by sonication using short but continuous pulses at 12 Hz for 1 min. This step was repeated 14 times to lyse all the cells. The lysate was centrifuged at 14000 rpm for 45 min at 4 °C to remove cell debris. The lysis suspension was brought to 30% saturation with ammonium sulfate and the pellet was discarded. This was followed by 50% saturation with ammonium sulfate. The solution was then centrifuged at 20000 rpm for 1 hour at 4 °C. The resultant pellet was dissolved in 10 mM Tris buffer, pH 7.4 and dialyzed overnight against the same buffer. After dialysis, the protein sample was filtered using a 30 kDa centricon filter (Merck Millipore, Darmstadt, Germany). The crude protein was then injected into a DEAE anion exchange column equilibrated with 10 mM Tris (pH 7.4) and eluted using a NaCl gradient. α-Syn was found to be eluted at about 300 mM NaCl. Fractions containing α-Syn (analyzed by Coomassie-stained SDS PAGE) were concentrated and further purified using a Sephadex gel filtration column. Fractions containing purified α-Syn were combined and lyophilized. The protein was determined to be about 95% pure by SDS-PAGE. For the sample preparations of all experiments, lyophilized protein was dissolved in sodium phosphate buffer (pH 7.4) and filtered using 0.22 μm low protein binding membranes (Millex-GP, Merck Millipore, Germany). The protein concentration was determined by the measurement of absorbance at 277 nm using the extinction coefficient 5960 cm^-1^M^-1^.

### Preparation of heme-treated oligomers_1_, oligomers_2_, untreated on-pathway oligomers and preformed seeds

For the preparation of the oligomers_1_ for cryo-EM, 200 μM lyophilized monomeric α-Syn was dissolved in sodium phosphate buffer, pH 7.4, and incubated for 20-24 h at 37 °C with constant agitation in the presence of 8 μM hemin chloride (pre-incubation). For the preparation of the untreated on-pathway oligomers, 200 μM monomeric α-Syn was incubated at 37 °C without agitation for 20-24 hours, after which it was ultracentrifuged at 45,000 rpm for 2 h and the supernatant was collected carefully while fibrils accumulated in the pellet were discarded. The excess monomeric species was removed by multiple filtration using a 100 kDa cutoff centricon (Amicon, Merck Millipore, MA, USA) to enrich the population of the oligomeric species. For the preparation of oligomers_2_, 200 μM prefibrillar and fibrillar aggregates (that had been previously incubated for 48 hours at 37 °C with constant agitation in the absence of hemin) were further incubated at 37 °C with constant agitation in presence of 8 μM hemin.

For the intracellular seeding experiments as well as Thioflavin T seeding assay, α-Syn seeds were produced by incubating 200 μM monomeric α-Syn protein for 96 hours at 37 °C under constant agitation, in the absence or presence of 8 μM hemin, after which the protein samples were sonicated in a bath sonicator for 15 minutes. For the Thioflavin T assay, 2 μM of the untreated or hemin-treated seeds were added to 200 μM fresh α-Syn monomers, and further incubated for a period of ∼ 72 hours.

### Thioflavin T assay

The formation of cross-β structure during the aggregation of α-Syn was measured by the addition of 20 μM ThT dissolved in buffer Thioflavin T (ThT) fluorescent probe to 2 μM protein aliquots collected from the incubation mixture at different time points (40). Changes in the emission fluorescence spectra recorded between 450 nm and 520 nm with the excitation wavelength set at 440 nm were monitored using a Photon Technology International fluorescence spectrometer.

### Negative Stain Electron Microscopy

5 μl aliquots taken from aggregation reactions were adsorbed onto 300 mesh carbon-coated copper grids (Agar Scientific, UK) and negative stained with 5 μl 2% (w/v) uranyl acetate. Images were obtained at various magnifications (1,000-90,000X) using a JEM-2100F 200 kV FE (Field Emission) transmission electron microscope.

### Atomic Force Microscopy

5 μl aliquots from the aggregation reactions were adsorbed onto a freshly cleaved muscovite mica (Agar Scientific, UK), followed by mild washing with 100 μl MilliQ water. The adsorbed α-Syn (treated or untreated with hemin) were imaged by Acoustic Alternative Current or AAC (tapping) mode using a Agilent Technologies Picoplus AFM 5500. The scan frequency was set at 1.5 Hz. A 9 µm scanner was used. All images were analyzed using Molecular Imaging Corporation PicoView 1.20.2 software (Molecular Imaging Corporation, CA, USA).

### Cell culture, α-Syn GFP transfection into SH-SY5Y neuroblastoma cells and treatment with untreated and heme-treated fibril seeds

The neuroblastoma cell line SH-SY5Y was maintained in DMEM supplemented with 10% heat-inactivated fetal bovine serum (FBS), 110 mg/L sodium pyruvate, 4 mM l-glutamine, 100 units/ml penicillin and 100 μg/ml streptomycin in humidified air containing 5% CO2 at 37 °C. The cells were transiently transfected with 2.5 μg wild type α-Syn-EGFP construct using Lipofectamine LTX and Plus reagent (Invitrogen, CA, USA) as described in the manufacturer’s protocol. 24 hours before the transfection, cells were seeded on 35 mm poly-D-lysine coated plates (MatTek Corporation, MA, USA) and allowed to grow till they were ∼ 60% confluent. For inducing the aggregation of the fusion protein, the cells were transduced using Lipofectamine with 2 μM fibril seeds dissolved in OptiMEM for 24 hours, after which they were washed twice with Dulbecco’s phosphate buffered saline (DPBS) and subjected to confocal imaging. α-Syn fibril seeds were prepared from the recombinant protein subjected to 96 h of aggregation, after which it was sonicated in a water bath for 15 minutes.

### Confocal Microscopy

These experiments were carried out using a Zeiss LSM 510 Meta confocal microscope equipped with a C-Apochromat 40 X (NA=1.20, water immersion) objective and confocal images were acquired with 512 × 512 (pinhole aperture ∼ 1 airy units). The α-Syn EGFP protein was excited using an argon laser at 488 nm.

### Preparation of SUVs (small unilamellar vesicles)

Monomeric and oligomeric α-Syn interact with acidic phospholipids, though the effects of oligomers on the dynamic properties of synthetic lipid vesicles also depend on the additional presence of neutral phospholipids.(41, 42) SUVs (small unilamellar vesicles) were used as α-Syn oligomers show a strong binding affinity to the augmented curvature of SUVs compared to that of LUVs (large unilamellar vesicles) and GUVs (giant unilamellar vesicles).(41) Due to its presence in brain membranes, phosphatidylserine (PS) was chosen instead of phosphatidylglycerol (PG) to increase the negative charge content of the vesicles.(41) Calcein-loaded SUVs of the composition of 3: 7 POPC: DOPS (POPC: 1-pamitoy-2-oleoyl-sn-glycero-3-phosphocholine; DOPS: 1, 2-dioleoyl-sn-glycero-3-phospho-L-serine) were added to α-Syn at a protein: lipid ratio of 1:10. 50 mM calcein when encapsulated within the SUVs is self-quenched, and hence shows a basal fluorescence at 515 nm when excited at 490 nm. 1 μl Triton X-100 was used to determine 100% calcein release, and all results were normalized to this value. For vesicle formation, a 3:7 ratio of POPC: DOPS was dissolved in 1 ml chloroform, followed by evaporation of the solvent under a stream of N_2_ gas. The resulting lipid film was hydrated in 20 mM sodium phosphate buffer, pH 7.4 containing 50 mM calcein dyes. The lipid-calcein suspension was sonicated in a glass tube in the dark at 40% amplitude for 30 minutes with 30 second pulse on and 1 minute pulse off at room temperature until the sample was transparent yellow in colour. The SUVs were isolated from free the dye by dialysing them in the dark in phosphate buffer, pH 7.4. The SUVs had a hydrodynamic diameter of 35±10 nm according to dynamic light scattering measurements. The hydrodynamic radius estimation was done using Malvern particle size analyser (Model no. ZEN 3690 ZETASIZER NANO ZS 90).

### Cytotoxicity assays

For the FITC-Annexin V/ Propidium Iodide early and late apoptosis assays, cells were seeded in 6 wells plates at 1×10^6^ cells/well. 24 hours after seeding, the cells were subjected to untreated α-Syn oligomers or oligomers_1_ or oligomers_2_ treated with heme at various time-points. For comparison, a sample with equivalent heme concentration was used as the control. The final concentration of the protein added to the cells was maintained at 10 μM. After addition of the treatments, the cells were incubated for 24 hours at 37 °C, 5% CO_2_. The percentage of apoptotic cells was determined using the Dead Cell Apoptosis Kit following instructions from the manufacturer. Apoptosis is a cellular process that entails a genetically programmed series of events leading to the death of a cell. During early apoptosis, the lipid phosphatidylserine (PS) is translocated to the outer side of the plasma membrane from the cytoplasmic side. FITC-conjugated Annexin V is a strong probe for the exposed PS and can thus be used for detecting early apoptosis in stressed cells.(43)

For the determination of late apoptosis and/or necrosis as a result of oligomer treatment, propidium iodide (PI) was added to the treated cells at a concentration of 2 μg/mL. PI labels the cellular DNA in late apoptotic/ necrotic cells where the cell membrane has been totally compromised.

### FTIR Analysis

FTIR measurements of the protein samples were performed in D_2_O-containing sodium phosphate buffer, pH 7.5 on a Bruker FTIR TENSOR 27 spectrometer. For the experiments, in order to study the effect of the heme on the *de novo* aggregation of α-Syn, 200 μM of the monomeric protein was incubated without or with 8 μM hemin under aggregation-inducing conditions (37°C, 180 rpm) for 20-24 hours. Conversely, to study the disaggregation of mature fibrils, another set of reactions was set up in which 200 μM mature fibrils (that had been incubated for 96 hours without hemin) were further incubated with 8 μM hemin for 48 hours. The protein concentration for each reading was maintained at 100 μM. The deconvoluted FTIR spectra of proteins in the Amide I region (1,700–1,600 cm^-1^) is due to the C=O stretching vibrations of the peptide bonds. It is predominantly informative about the backbone conformation and the relative composition of secondary structure elements of the protein (27). The buffer background was independently measured and subtracted from each protein spectrum before curve fitting of the Amide I region. Data presented are a result of the fitting of 3 independent FTIR spectra for each protein sample. In order to estimate the relative fraction of β-sheet content in each protein sample, deconvolution analysis with Gaussian/Lorentzian curves for each spectra recorded was performed. The different Gaussian distributions were consigned to contributions from either β-sheet secondary structure or turns, or random coil according to the position of their peaks (44). The analysis of the FT-IR spectra allows the distinction between parallel and anti-parallel β-sheet structures based on the analysis of the amide I (1700-1600 cm^-1^) region (45). In anti-parallel β-sheet structures, the amide I region displays two typical components: the major low frequency one has an average wavenumber located at ∼1620-1638 cm^-1^, whereas the minor high frequency component, 4-5-fold weaker than the major one, is characterized by an average wavenumber at 1695 cm^-1^. For parallel β-sheet structures, the amide I region displays only the major component around 1620-1638 cm^-1^. The oligomers and the fibrils reflect a high resemblance in the type of secondary structure organization except that they differ in the extent of β-sheet versus disordered content and the overall β-sheet arrangement (parallel for the fibrillar state and anti-parallel for the oligomeric state). Turns are associated with various bands between 1660 and 1690 cm^-1^, while unordered regions and loops are represented by bands around ∼1642-46 cm^-1^ and ∼1656-64 cm^-1^, respectively. Solvent subtraction, self-Fourier deconvolution of the Amide I region, determination of band position and curve-fitting were performed using OriginPro 8.5 software (OriginLab Corp., MA, USA).

### Hemin-α-Syn Binding

In order to study the binding of hemin to α-Syn, we incorporated a single cysteine mutation at the C-terminal of α-Syn to which a maleimide-containing dye could be attached. The binding assays of TMR-5-maleimide tagged α-Syn G132C or H50Q/G132C was investigated at 25 °C by fluorescence spectroscopy. The tagging of the protein using the dye was performed following the manufacturer’s protocol. The measurements were performed on a Photon Technology International fluorescence spectrometer using a 1.0 cm path-length cell. Titrations were performed by adding increasing concentrations (0.1 μM - 4 μM) of hemin dissolved in sodium phosphate buffer, pH 7.4, to a fixed amount (0.1 μM) of α-Syn G132C or H50Q/G132C tagged with Tetramethylrhodamine-5-maleimide (ex/em 550nm/ 574nm in sodium phosphate buffer). The quenching efficiency was then evaluated by the Stern-Volmer quenching constant (K_SV_), which is calculated from the following equation:(46)

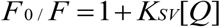

where F_0_ and F are the emission intensities of α-Syn-TMR-5-maleimide in absence and presence of different concentrations of heme, and [Q] is the concentration of heme (quencher). A plot of F_0_/F versus [Q] yields a slope equal to the Stern-Volmer quenching constant (K_SV_). The relationship between the fluorescence intensity of the dye-tagged protein and the concentration of quencher was utilised to obtain the association constant (K_a_) and the number of binding sites (n), both of which were calculated from the following equation:

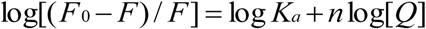

where, K_a_ refers to the association constant. The dissociation constant K_d_ = 1/ K_a_.

### Determination of Peroxidase Activity

For the estimation of the peroxidase activity of α-Syn in the absence and presence of heme, 2 μM heme-bound or unbound α-Syn (WT or H50Q, α-Syn/hemin 25:1) was treated at 25°C with 200 µM H_2_O_2_ and 10 mM guaiacol. The product formation kinetics was followed using a Shimadzu 1700 Pharmaspec UV-VIS spectrophotometer. The rate of decomposition of hydrogen peroxide (H_2_O_2_) by peroxidase using guaiacol as a hydrogen donor, was determined by measuring the rate of color development spectrophotometrically at 470 nm using the extinction coefficient of 2.66X10^4^ M^-1^ cm^-1^ (47).

### FPLC-SEC

The FPLC-size exclusion chromatography analysis was performed by injecting samples of heme-treated oligomers_1_ and oligomers_2_ onto a Bio-Rad ENrich SEC70 10 x 300 mm column using the Bio-Rad NGC FPLC system (Bio-Rad, CA, USA). The retention time of both oligomer types was compared with a set of molecular weight protein standards (Bio-Rad) consisting of bovine serum albumin (66.5 kDa), ovalbumin (44 kDa), myoglobin (17 kDa) and vitamin B12 (1.35 kDa). All FPLC runs were performed in pH 7.4 sodium phosphate buffer at a flow rate of 0.5 ml/min and monitored by absorbance at 280 nm.

### Cryo-electron microscopy

Samples (∼5 µl) were applied on glow-discharged lacey grids (Ted Pella, CA, USA), followed by blotting and vitrification of the grids using Vitrobot(tm) (FEI, OR, USA) (48). Image data collection was performed on a Tecnai POLARA microscope (FEI, OR, USA) equipped with a FEG (Field Emission Gun) operating at 300 kV. Images were collected with 4K X 4K ‘Eagle’ charge-coupled device (CCD) camera (FEI, OR, USA) at ∼79000X magnification (with defocus values ranging from ∼1 to 5 μm), resulting in a pixel size of 1.89 Å at the object scale. All images were acquired using low-dose procedures with an estimated dose of ∼20 electrons per Å_2_ (49). Micrograph screening and particle picking were done separately using EMAN2 (30) and SPIDER (50). 60 micrographs for Sample 1 (α-Syn pre-incubated with heme for 24 h), 41 micrographs for Sample 2 (α-Syn late-incubated with heme after 48 h of aggregation), and 118 micrographs for Sample 3 (α-Syn incubated without heme for 24 h) were selected for particle picking. The initial models for the three different samples were developed using EMAN2.

Validation of the 3D models was done in multiple ways. Particles formed following heme treatment appeared to be very small (∼5-8 nm). We imaged three sets of data for each of the samples (each prepared and imaged on different dates) to confirm the sizes of the particles, particularly for oligomers pre-incubated with heme (sample 1), which showed distribution of particles of small sizes. The different views of the reference-free 2D class averages generated by different image processing softwares, e.g., EMAN2, Xmipp (51), RELION (52) as well as reference-based SPIDER were very similar. The initial models generated in EMAN2 were used as references for SPIDER auto picking after which 17831, 18626 and 11515 good particles were selected manually from auto-picked particles of Samples 1, 2 and 3, respectively. The 3D reconstruction was then performed following the standard SPIDER protocol for reference-based reconstruction (50) for each dataset. The 2D re-projections of 3D maps were consistent with the 2D class averages. The overall resolution of three maps generated for samples 1, 2, and 3 were 12.6 Å, 12.0 Å and 16.0 Å, respectively, using the FSC 0.5 cutoff criteria (8.9 Å, 8.9 Å and 11.0 Å using the 0.143 cutoff criteria, Figure S2). We also manually produced another starting model from the Greek key structural motif (as a tetramer) by low pass filtering. The density model showed no distortion at the head and base junction. 2D back projections of this 3D structure also resembled 2D class averages of samples 1 and 2 (Figure S2). Final 3D map generated from dataset of sample 1 (oligomers_1_) using this initial model showed similar ‘molecular mace’ architecture with a twist at the head and base junction.

Surface rendering, docking of crystal structures, segmentation and analyses of the 3D maps were performed in the program UCSF Chimera (53), Pymol (54), and VMD (55).

## Acknowledgement

This work was supported by CSIR Network project ‘UNSEEN’ (BSC0113). We greatly appreciate the help from the Central Instrumentation Facilities (CIF) and the technical persons attached to it at the CSIR-Indian Institute of Chemical Biology. We sincerely thank Mr. Chiranjit Biswas for the cryo-EM data collection of the protein-heme complexes. We acknowledge Mr Thangamuniyandi Muruganandan, CIF, CSIR-IICB and Mr Supriya Chakraborty, CSS, Indian Association for the Cultivation of Science, for their assistance with the AFM and TEM imaging and analyses. RC, SD and PS acknowledge the UGC for their senior research fellowship; SSP expresses gratitude to the CSIR for his fellowship. We thank the Director, CSIR-IICB for his help and encouragements.

## Author contributions

KC conceived the project. KC, RC and SSP designed the experiments in consultation with JS for the cryo-EM section. RC carried out the biochemical and cell-based experiments. SSP and PS performed initial biochemical experiments and image processing of the cryo-EM data. SD assisted in few biochemical experiments and performed elaborate 3D cryo-EM image processing and validation of the maps. KC, JS, RC and SD analyzed the data and wrote the paper.

## References

1. Spillantini MG, et al. (1997) Alpha-synuclein in Lewy bodies. Nature 388(6645):839-840.

2. Dunnett SB & Bjorklund A (1999) Prospects for new restorative and neuroprotective treatments in Parkinson’s disease. Nature 399(6738 Suppl):A32–39.

3. Iljina M, et al. (2016) Kinetic model of the aggregation of alpha-synuclein provides insights into prion-like spreading. Proceedings of the National Academy of Sciences 113(9):E1206–E1215.

4. Garcia GA, Cohen SIA, Dobson CM, & Knowles TPJ (2014) Nucleation-conversion-polymerization reactions of biological macromolecules with prenucleation clusters. Physical Review E 89(3):032712.

5. Winner B, et al. (2011) In vivo demonstration that alpha-synuclein oligomers are toxic. Proceedings of the National Academy of Sciences of the United States of America 108(10):4194–4199.

6. Lashuel HA, et al. (2002) Alpha-synuclein, especially the Parkinson’s disease-associated mutants, forms pore-like annular and tubular protofibrils. Journal of molecular biology 322(5):1089–1102.

7. Uversky VN (2008) Amyloidogenesis of natively unfolded proteins. Curr Alzheimer Res 5(3):260–287.

8. Tuttle MD, et al. (2016) Solid-state NMR structure of a pathogenic fibril of full-length human alpha-synuclein. Nature structural & molecular biology 23(5):409–415.

9. Li B, et al. (2018) Cryo-EM of full-length α-synuclein reveals fibril polymorphs with a common structural kernel. Nature communications 9(1):3609–3609.

10. Guerrero-Ferreira R, et al. (2018) Cryo-EM structure of alpha-synuclein fibrils. eLife 7:e36402.

11. Li Y, et al. (2018) Amyloid fibril structure of alpha-synuclein determined by cryoelectron microscopy. 28(9):897–903.

12. Kayed R, et al. (2003) Common structure of soluble amyloid oligomers implies common mechanism of pathogenesis. Science 300(5618):486–489.

13. Chen SW, et al. (2015) Structural characterization of toxic oligomers that are kinetically trapped during α-synuclein fibril formation. Proceedings of the National Academy of Sciences 112(16):E1994–E2003.

14. Fusco G, et al. (2017) Structural basis of membrane disruption and cellular toxicity by alpha-synuclein oligomers. Science 358(6369):1440–1443.

15. Hayden EY, et al. (2015) Heme Stabilization of α-Synuclein Oligomers during Amyloid Fibril Formation. Biochemistry 54(30):4599–4610.

16. Lamberto GR, et al. (2011) Toward the Discovery of Effective Polycyclic Inhibitors of α-Synuclein Amyloid Assembly. The Journal of Biological Chemistry 286(37):32036–32044.

17. Fonseca-Ornelas L, et al. (2014) Small molecule-mediated stabilization of vesicle-associated helical alpha-synuclein inhibits pathogenic misfolding and aggregation. Nature communications 5:5857.

18. Chakraborty R, Sahoo S, Halder N, Rath H, & Chattopadhyay K (2018) Conformational-Switch Based Strategy Triggered by [18] π Heteroannulenes toward Reduction of Alpha Synuclein Oligomer Toxicity. ACS Chemical Neuroscience.

19. Yang W, Li X, Li X, Li X, & Yu S (2016) Neuronal hemoglobin in mitochondria is reduced by forming a complex with α-synuclein in aging monkey brains. Oncotarget 7(7):7441–7454.

20. Kleinknecht A, et al. (2016) C-Terminal Tyrosine Residue Modifications Modulate the Protective Phosphorylation of Serine 129 of α-Synuclein in a Yeast Model of Parkinson’s Disease. PLoS genetics 12(6):e1006098–e1006098.

21. Campioni S, et al. (2010) A causative link between the structure of aberrant protein oligomers and their toxicity. Nature chemical biology 6(2):140–147.

22. Zampagni M, et al. (2011) A comparison of the biochemical modifications caused by toxic and non-toxic protein oligomers in cells. Journal of cellular and molecular medicine 15(10):2106–2116.

23. Basak S, Prasad GVRK, Varkey J, & Chattopadhyay K (2015) Early Sodium Dodecyl Sulfate Induced Collapse of α-Synuclein Correlates with Its Amyloid Formation. ACS Chemical Neuroscience 6(2):239–246.

24. Joshi N, et al. (2015) Attenuation of the Early Events of α-Synuclein Aggregation: A Fluorescence Correlation Spectroscopy and Laser Scanning Microscopy Study in the Presence of Surface-Coated Fe3O4 Nanoparticles. Langmuir 31(4):1469–1478.

25. Angot E, et al. (2012) Alpha-Synuclein Cell-to-Cell Transfer and Seeding in Grafted Dopaminergic Neurons In Vivo. PLOS ONE 7(6):e39465.

26. Luk KC, et al. (2009) Exogenous α-synuclein fibrils seed the formation of Lewy body-like intracellular inclusions in cultured cells. Proceedings of the National Academy of Sciences 106(47):20051–20056.

27. Arrondo JL, Muga A, Castresana J, & Goni FM (1993) Quantitative studies of the structure of proteins in solution by Fourier-transform infrared spectroscopy. Progress in biophysics and molecular biology 59(1):23–56.

28. Celej MS, et al. (2012) Toxic prefibrillar alpha-synuclein amyloid oligomers adopt a distinctive antiparallel beta-sheet structure. The Biochemical journal 443(3):719–726.

29. Frank J (2002) Single-particle imaging of macromolecules by cryo-electron microscopy. Annu Rev Biophys Biomol Struct 31:303–319.

30. Tang G, et al. (2007) EMAN2: an extensible image processing suite for electron microscopy. Journal of structural biology 157(1):38–46.

31. Shaikh TR, et al. (2008) SPIDER image processing for single-particle reconstruction of biological macromolecules from electron micrographs. Nature protocols 3(12):1941–1974.

32. Quist A, et al. (2005) Amyloid ion channels: a common structural link for protein-misfolding disease. Proceedings of the National Academy of Sciences of the United States of America 102(30):10427–10432.

33. Wu K-P, Kim S, Fela DA, & Baum J (2008) Characterization of conformational and dynamic properties of natively unfolded human and mouse alpha-synuclein ensembles by NMR: implication for aggregation. Journal of molecular biology 378(5):1104–1115.

34. Pramanik D & Dey SG (2011) Active Site Environment of Heme-Bound Amyloid β Peptide Associated with Alzheimer’s Disease. Journal of the American Chemical Society 133(1):81–87.

35. Appel-Cresswell S, et al. (2013) Alpha-synuclein p.H50Q, a novel pathogenic mutation for Parkinson’s disease. Movement Disorders 28(6):811–813.

36. Trabuco LG, Villa E, Schreiner E, Harrison CB, & Schulten K (2009) Molecular dynamics flexible fitting: a practical guide to combine cryo-electron microscopy and X-ray crystallography. Methods 49(2):174–180.

37. Chakraborty R & Chattopadhyay K (2019) Cryo-Electron Microscopy Uncovers Key Residues within the Core of Alpha-Synuclein Fibrils. ACS Chemical Neuroscience.

38. Bartels T, Choi JG, & Selkoe DJ (2011) alpha-Synuclein occurs physiologically as a helically folded tetramer that resists aggregation. Nature 477(7362):107–110.

39. Wang W, et al. (2011) A soluble α-synuclein construct forms a dynamic tetramer. Proceedings of the National Academy of Sciences 108(43):17797–17802.

40. LeVine H, 3rd (1999) Quantification of beta-sheet amyloid fibril structures with thioflavin T. Methods in enzymology 309:274–284.

41. van Rooijen BD, Claessens MM, & Subramaniam V (2009) Lipid bilayer disruption by oligomeric alpha-synuclein depends on bilayer charge and accessibility of the hydrophobic core. Biochimica et biophysica acta 1788(6):1271–1278.

42. Volles MJ, et al. (2001) Vesicle permeabilization by protofibrillar alpha-synuclein: implications for the pathogenesis and treatment of Parkinson’s disease. Biochemistry 40(26):7812–7819.

43. Engeland Mv, Nieland LJW, Ramaekers FCS, Schutte B, & Reutelingsperger CPM (1998) Annexin V-Affinity assay: A review on an apoptosis detection system based on phosphatidylserine exposure. Cytometry 31(1):1–9.

44. Anonymous (Infrared Spectroscopy of Proteins. Handbook of Vibrational Spectroscopy..

45. Cerf E, et al. (2009) Antiparallel beta-sheet: a signature structure of the oligomeric amyloid beta-peptide. The Biochemical journal 421(3):415–423.

46. Lakowicz JR (1999) Principles of fluorescence spectroscopy (Second edition. New York: Kluwer Academic/Plenum, [1999] ©1999).

47. Diederix RE, Ubbink M, & Canters GW (2002) Peroxidase activity as a tool for studying the folding of c-type cytochromes. Biochemistry 41(43):13067–13077.

48. Grassucci RA, Taylor DJ, & Frank J (2007) Preparation of macromolecular complexes for cryo-electron microscopy. Nature protocols 2(12):3239–3246.

49. Grassucci RA, Taylor D, & Frank J (2008) Visualization of macromolecular complexes using cryo-electron microscopy with FEI Tecnai transmission electron microscopes. Nature protocols 3(2):330–339.

50. Shaikh TR, et al. (2008) SPIDER image processing for single-particle reconstruction of biological macromolecules from electron micrographs. Nature Protocols 3(12):1941–1974.

51. Sorzano COS, et al. (2004) XMIPP: a new generation of an open-source image processing package for electron microscopy. Journal of Structural Biology 148(2):194–204.

52. Scheres SHW (2012) RELION: Implementation of a Bayesian approach to cryo-EM structure determination. Journal of Structural Biology 180(3):519–530.

53. Pettersen EF, et al. (2004) UCSF chimera - A visualization system for exploratory research and analysis. Journal of Computational Chemistry 25(13):1605–1612.

54. The PyMOL Molecular Graphics System DSL, San Carlos, CA. (2002).

55. Humphrey W, Dalke, A. and Schulten, K., (1996) VMD - Visual Molecular Dynamics. J. Molec. Graphics 14:33–38.

