## Supporting Information for "Heme minimizes Parkinson’s disease-associated toxicity by inducing a conformational distortion in the oligomers of alpha-Synuclein"

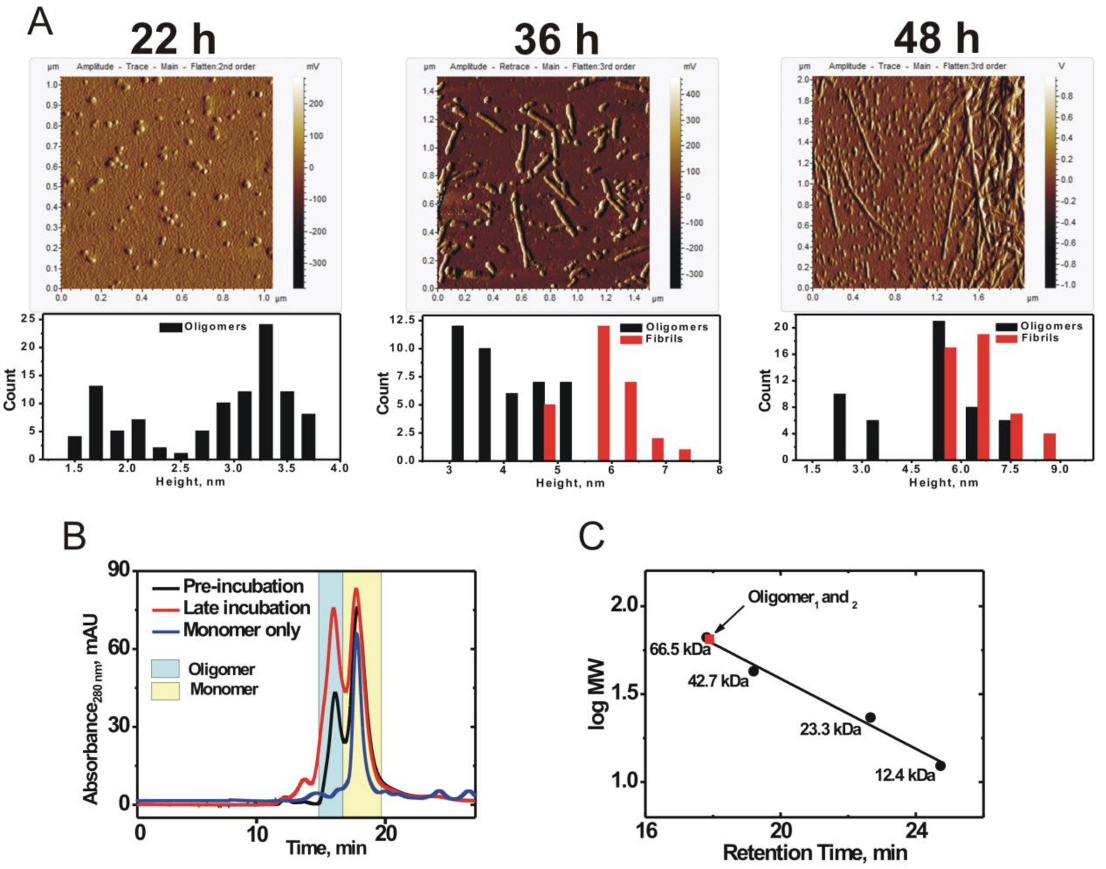


**Figure S1:** (A) AFM micrographs and their corresponding size distribution profile of the aggregates formed between 24 h and 48h. (B) FPLC-SEC chromatogram depicting the formation of the mace oligomers when heme is added to α-Syn at an early or late occasion. In both cases, the oligomers$_{1\text{ and }2}$ have an identical retention time of 17.1 minutes, while the monomeric population elutes at 18.2 minutes. (C) The retention time for each protein standard used for the calibration are shown and compared with the heme-stabilized oligomer and the extended (13) monomer.

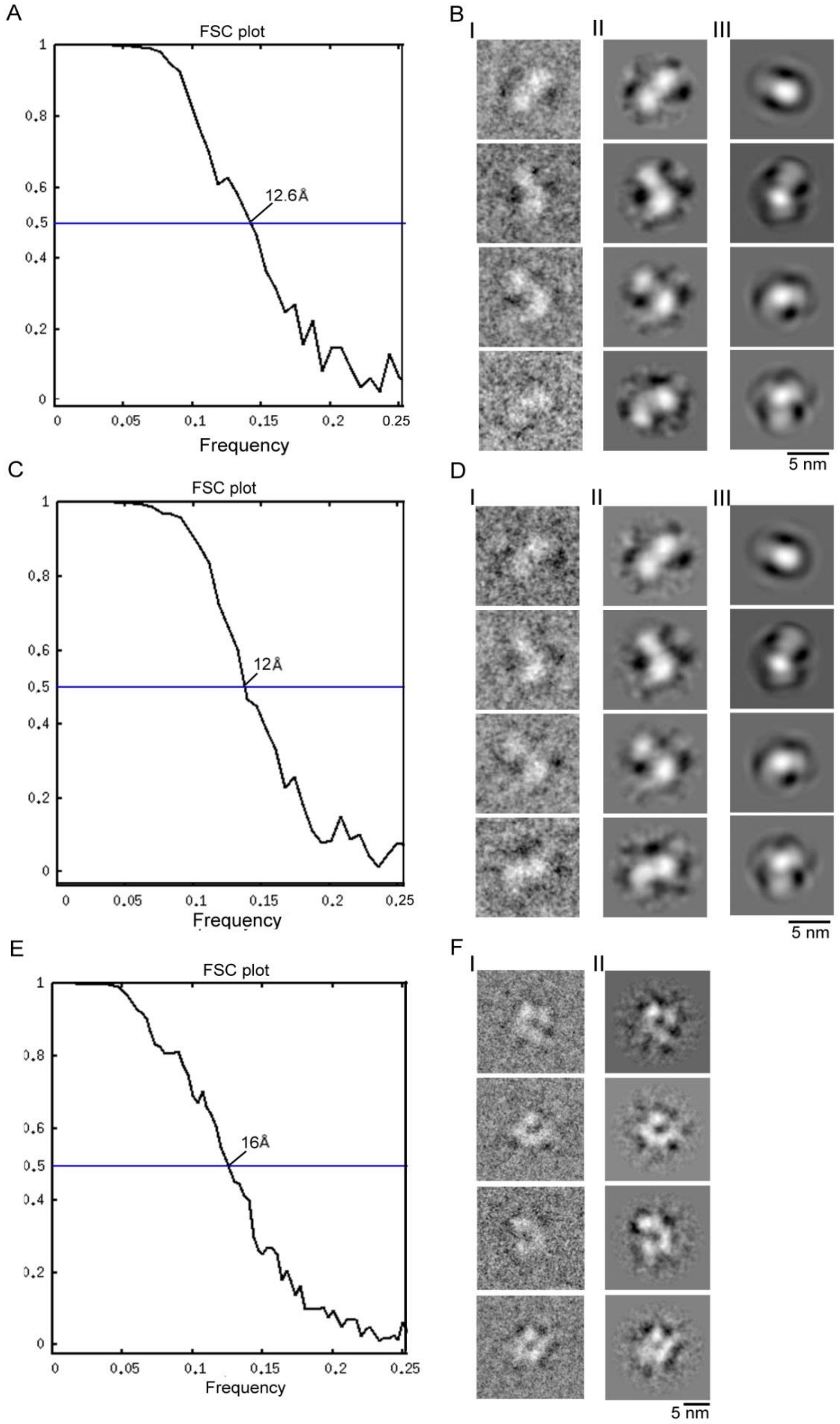


**Figure S2: Analysis of the cryo-EM image processing of three datasets.** FSC plots (A, C and E) show resolutions 12.6Å, 12Å and 16Å of sample 1, the ‘molecular mace’ oligomer when heme treated at the begining ($oligomers_1$) (A); sample 2, ‘molecular mace’ oligomer when heme treated after fibril formation ($oligomers_2$) (C); and sample 3, horseshoe oligomer formed in the absence of heme (E), respectively, at 0.5 cutoff of the FSC curve. (B, D, and F) show comparison of 2D averages for samples 1, 2, and 3, respectively. Different views (I) of 2D class averages of particles and projection of final maps (II) are compared. 2D projections of density map created from a taken 2N0A PDB structure (as tetramer) are shown in III (B, D).

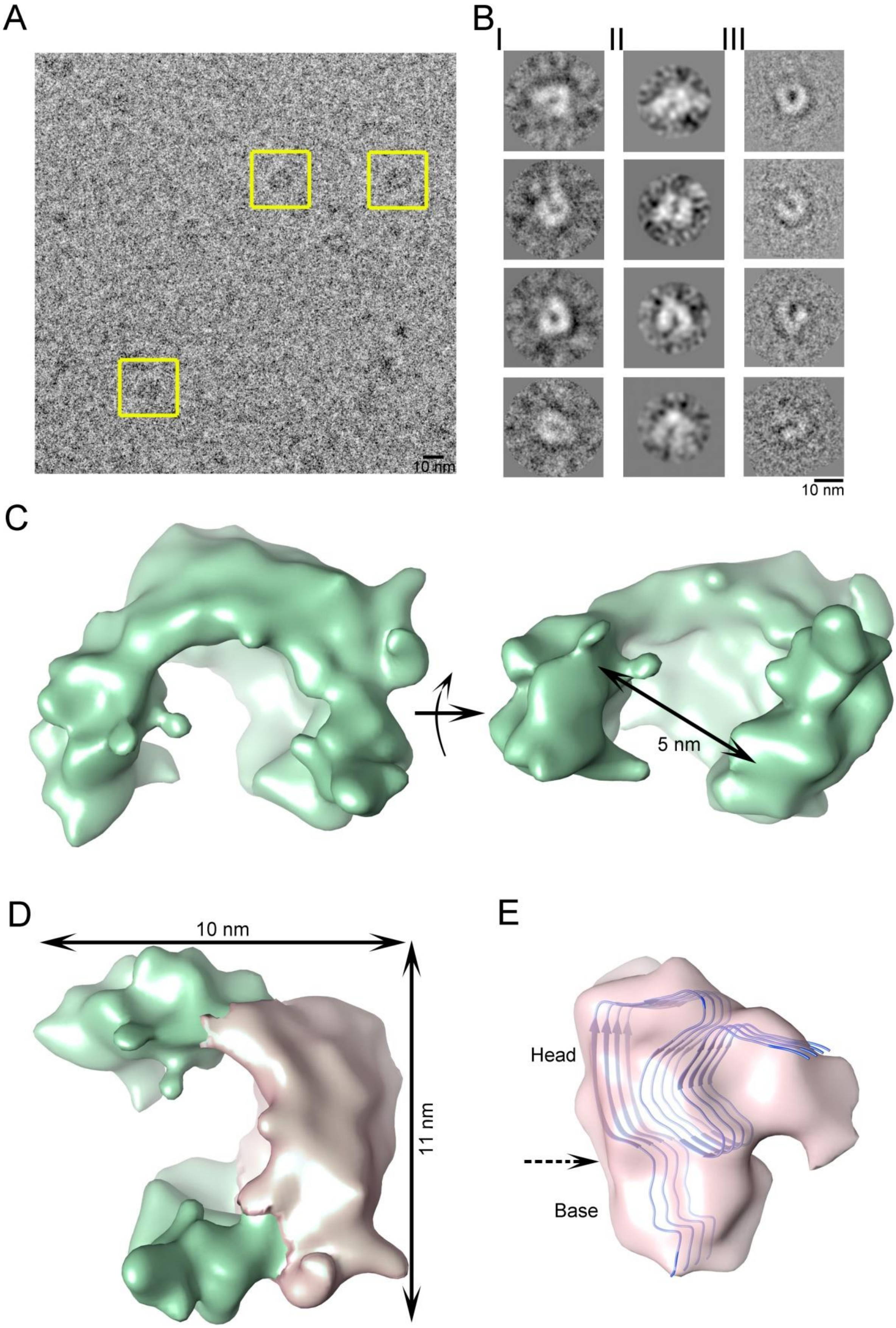

A
10 nm
B
I
II
III
10 nm
C
5 nm
D
10 nm
11 nm
E
Head
Base

**Figure S3**: Cryo-EM study of (A-C) the oligomers formed ~24 h of aggregation in the absence of heme. (A) Micrograph depicting distribution of a prominant oligomeric species in a heterogeneous population (yellow box). (B I-III) Reference free 2D class averages generated in Xmipp, RELION and EMAN2, respectively, showing horseshoe shape of the oligomer. (C) Two views of the 3D cryo-EM density map of the horseshoe oligomer of α-Syn formed in the absence of heme. Apparently, this form might be a precursor of previously reported annular oligomeric forms. (D) One of the segment of the density map is overlaid (pink, segmentation done using Chimera). (E) Docking of α-Syn Greek key motif (tetrameric unit, blue cartoon) into isolated density segment (semitransparent pink) shows accomodation of the coordinates without any requirement of distortion at the junction of 'head' and 'base' (marked by dotted arrow) indicating that tetrameric greek key motif is likely the fundamental unit for α-Syn oligomerization.

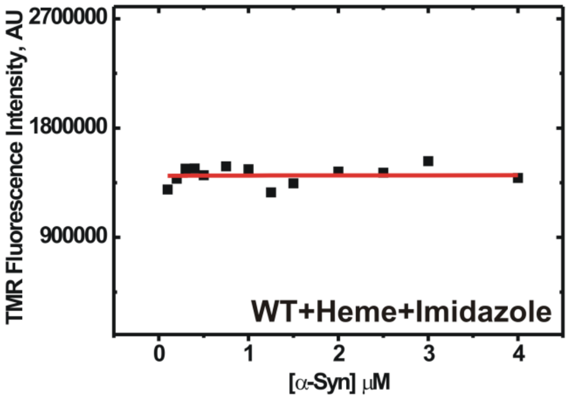


**Figure S4:** Heme shows no binding to WT α-Syn shows in presence of excess imidazole (10 mM) which competes with the histidine residue of the protein.
